# Photoaged microplastics disrupt endothelial stretch-sensitive ion channels to impair calcium signaling and vascular integrity

**DOI:** 10.64898/2026.05.01.722241

**Authors:** Seul-Ki Park, Jae Min Cho, Enbo Zhu, Khoa Vu, Jing Wang, Peng Zhao, Aaron S. Romero, Matthew J. Campen, Srinivasa T. Reddy, Eliseo F. Castillo, Tzung Hsiai

**Affiliations:** Division of Cardiology, Department of Medicine, David Geffen School of Medicine, University of California, Los Angeles, CA, 90095, USA; Department of Medicine, Greater Los Angeles Veteran Affairs Healthcare System, Los Angeles, CA, 90095, USA; Department of Molecular and Medical Pharmacology, David Geffen School of Medicine, University of California, Los Angeles, CA, 90095, USA; Division of Gastroenterology and Hepatology, Department of Internal Medicine, University of New Mexico Health Sciences Center, Albuquerque, NM, 87106, USA; Department of Pharmaceutical Sciences, College of Pharmacy, University of New Mexico Health Sciences, Alburquerque, NM, 87106, USA

## Abstract

Plastic-derived micro- and nanoplastics are pervasive, but how environmentally aged particles affect vascular barriers is poorly understood. We hypothesized that photoaged plastics impair endothelial force-sensing, triggering gut-brain-heart barrier failure. Ultraviolet (UV) exposure converted pristine nanoplastics into oxidized, irregular photoaged microplastic aggregates (> 1.2 µm). In human aortic endothelial cells, photoaged particles increased membrane stiffness and activated transcriptional programs linked to permeability, junction disruption, inflammation, and cytoskeletal remodeling. Mechanistically, photoaged particles selectively inhibited Piezo1-mediated Ca^2+^ signaling and downstream Notch activity without changing PIEZO1 expression, and endothelial CRISPR inhibition of PIEZO1 recapitulated these effects. In zebrafish, photoaged plastic exposure increased gut-vascular permeability and systemic spread with brain and heart accumulation, accompanied by reduced neurovascular and myocardial Ca^2+^ signals, depressed cardiac contractility, and abnormal locomotor behavior. Thus, photoaged plastics compromise vascular barriers through disrupted endothelial Piezo1-Notch mechanotransduction.

## Introduction

Microplastics (<5 mm, MPs) and nanoplastics (<1 µm, NPs) (NMPs), collectively referred to as nano- and microplastics (NMPs), have emerged as pervasive environmental contaminants that accumulate across natural and human associated ecosystems (*1, 2*). Although some NMPs are released directly from primary sources such as microbeads, the majority arise from the photooxidative weathering of larger plastic debris under environmental stressors including ultraviolet (UV) radiation, thermal cycling, and oxidative conditions (*1, 3*). These processes generate structurally and chemically heterogeneous particles with altered surface chemistry, irregular morphologies, and increased reactivity (*4*). UV-mediated aging, often accompanied by microbial colonization, accelerates surface oxidation and enhances biological interactions, thereby increasing particle bioavailability and toxicological potential (*5, 6*). Consistent with this, environmental plastics have been detected in human blood (*7, 8*), lungs (*9*), liver (*10*), placenta (*11*), and heart (*12*), and postmortem brain tissues (*13*), raising concerns that vascular barriers may be particularly vulnerable to exposure.

The plasma membrane represents a critical interface at which environmental particles first interact with cells, regulating ion flux, metabolite exchange, and mechanochemical signaling essential for cellular homeostasis (*14, 15*). In endothelial cells, calcium signaling plays a central role in maintaining barrier integrity by coordinating tight junction dynamics, angiogenesis, and redox balance (*16*). The mechanosensitive ion channel PIEZO1, activated by shear stress, mediates calcium influx (*17*) and engages downstream pathways including Notch signaling, a key regulator of vascular patterning, endothelial stability, and anti-inflammatory responses (*18-20*). Although Piezo1-Notch signaling is essential for endothelial resilience, whether environmentally aged, photooxidized microplastics disrupt this mechanotransductive axis remains unknown.

Here, we investigate how photoaged microplastics (PA-MPs) influence endothelial mechanosensitive signaling and vascular barrier function. We show that PA-MPs impair Piezo1-dependent calcium entry, suppress downstream Notch signaling, and compromise endothelial barrier integrity, thereby promoting translocation across the gut-vascular interface and systemic distribution to distant organs. Together, this work establishes endothelial mechanotransduction as a critical interface through which environmentally aged plastics compromise vascular integrity and promote systemic distribution.

## Results

### PA-MPs induce oxidative stress and biomechanical remodeling in endothelial cells

The morphological characteristics of pristine nanoplastics (NPs) and photoaged microplastics (PA-MPs), generated by UV irradiation of NPs, were examined using scanning electron microscopy (SEM) (Fig. 1A). Because photoaging-induced aggregation and fragmentation of polystyrene particles are strongly influenced by surrounding ionic conditions(*21*), we compared the effects of UV exposure on NP morphology between E3 medium versus double-distilled water (DDW) to distinguish intrinsic photoaging from ion-dependent transformation. Following 2 weeks of UV exposure, 50 nm NPs exhibited the onset of surface roughening and a shift toward larger apparent particle size under both E3 medium and DDW conditions. At this time point, particles aged in E3 medium showed enhanced clustering and greater size heterogeneity compared with those aged in DDW. In contrast, prolonged UV irradiation for 4 weeks resulted in evident surface roughening, deformation, and increased structural heterogeneity under both conditions (Fig. 1A). Particles aged in E3 medium displayed distinct clustered size distributions, characterized by particle fragmentation, self-aggregation, and the emergence of irregular, porous surface features compared with DDW. Particles subjected to 4 weeks of UV irradiation are hereafter referred to as photoaged microplastics (PA-MPs).

**Fig. 1.**
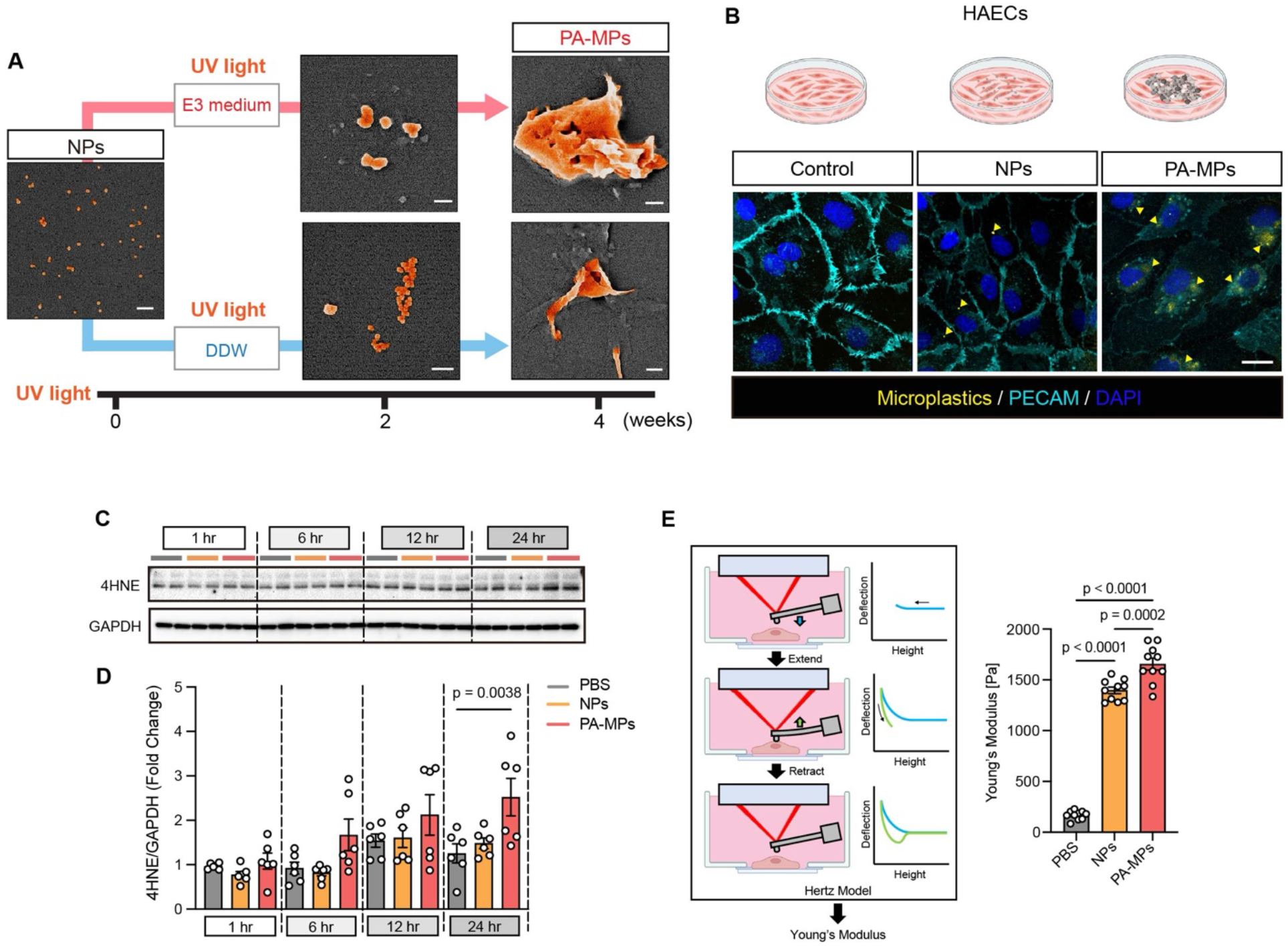
PA-MPs alter particle morphology, induce oxidative stress, stiffen endothelial cells, and impair mechanosensitive Ca^2+^ signaling. (**A**) Representative scanning electron microscopy (SEM) images of fresh polystyrene nanoplastics (PS-NPs; d = 50 nm) and photoaged microplastics (PA-MPs; d ≥ 1.2 µm) following ultraviolet (UV) irradiation for 2 or 4 weeks in E3 medium (upper panels) or double-distilled water (DDW; lower panels). Scale bar: 200 nm. (**B**) Immunofluorescence staining with a dye against microplastics (yellow) and PECAM-1 (cyan) shows intracellular microplastic localization within HAECs (yellow arrowheads: internalized microplastic particles). Scale bar: 20 µm. (**C** and **D**) Representative immunoblots showing progressive increases in oxidative stress, as indicated by elevated 4-hydroxynonenal (4-HNE) levels, with prolonged PA-MPs exposure relative to PBS controls and NPs-treated cells. (**E**) Atomic force microscopy (AFM) measurements demonstrate that PA-MPs exposure significantly increases endothelial cell stiffness, quantified by Young’s modulus (n = 10 cells per condition).

Firstly, to optimize the concentration of NPs and PA-MPs in endothelial cells, we evaluated the cytotoxic effects of NPs and PA-MPs in human aortic endothelial cells (HAECs) to determine appropriate exposure conditions for subsequent analyses. PA-MPs exhibited greater cytotoxic effects relative to pristine particles, consistent with previous reports on aged microplastics (*22*). Accordingly, an exposure condition corresponding to approximately a 30% reduction in cell viability for PA-MPs (8 ug/ml), compared with ∼12% for NPs, was selected to avoid overt cytotoxicity (fig. S1A). All subsequent experiments were conducted under this exposure condition. Additionally, EdU incorporation assays were performed to assess whether PA-MPs exposure affects the proliferative capacity alongside cytotoxicity. PA-MPs treatment resulted in a significant reduction in EdU-positive cells relative to controls, suggesting compromised endothelial proliferative potential (fig. S1, B and C).

We next confirmed the presence and intracellular localization of NPs and PA-MPs within HAECs using a particle-associated fluorescent signal detection reagent, a conjugated polymer nanoparticle-based fluorescent probe validated for reliable imaging in biological samples (Stable staining of microplastics using conjugated polymer nanoparticles (*23*). We observed that NPs exhibited a diffuse intracellular distribution, whereas PA-MPs were detected as larger, discernible aggregates, frequently localizing in the perinuclear region as encapsulated clusters within ECs. These particles displayed unique intracellular morphologies, consistent with the differential surface features revealed by SEM (Fig. 1A). Correspondingly, levels of 4-hydroxynonenal (4-HNE), a marker of lipid peroxidation and oxidative stress, progressively increased in PA-MPs-treated HAECs compared with PBS or NPs treated controls (Fig. 1, C and D). To test endothelial biomechanical properties, we measured Young’s Modulus, an indicator of cell stiffness, using atomic force microscopy (AFM). Exposure to both NPs and PA-MPs robustly increased EC stiffness relative to PBS-treated HAECs; however, PA-MPs-treated cells exhibited higher stiffness than NPs-treated cells (Fig. 1E).

### PA-MPs compromise mechanosensitive ion channels in endothelial cells

We further explored mechanosensitive ion channel function in response to PA-MPs. Intracellular Ca^2+^ dynamics were quantified as relative fluorescence intensity (ΔF/F_0_) using a calcium-sensitive indicator. In the control condition, Yoda1, a Piezo1 specific agonist that stabilizes the channel in an open, mechanically active conformation (*24-26*) elicited a rapid transient increase in intracellular Ca^2+^, followed by a gradual decay toward baseline levels, a response that was preserved in HAECs treated with NPs, however, HAECs exposed to PA-MPs exhibited a significantly attenuated Ca^2+^ in response to Yoda1 (Fig. 2, A to C and movie S1). Pharmacological inhibition of mechanosensitive ion channels with GsMTx4, a non-selective mechanosensitive ion channel inhibitor, suppressed Yoda1-induced Ca^2+^ influx, recapitulating the effects observed with PA-MPs exposure (Fig. 2, A and B). To assess whether other mechanosensitive ion channels are involved in PA-MPs-induced Ca^2+^ influx, TRPV4 and TRPC6 channels were selectively activated using the agonists GSK1016790A and Hyp9, respectively. Both TRPV4 and TPRC6 dependent Ca^2+^ influx was reduced by PA-MPs (Fig. 2, D to F), suggesting a generalized impairment of mechanosensitive Ca^2+^ entry rather than a selective inhibition of PIEZO1. Quantitative analysis of maximal Ca^2+^ responses, expressed as peak ΔFmax/F_0_, showed that activation of PIEZO1 with Yoda1 elicited higher-amplitude Ca^2+^ signals than stimulation of TRPV4 or TRPC6 (Fig. 2, C and F). PA-MPs exposure reduced maximal Ca^2+^ entry across all mechanosensitive channels examined, whereas PIEZO1-mediated responses exhibited the greater reduction in peak amplitude compared to TRPV4 and TRPC6 indicating that Piezo1 is a major regulator of PA-MPs induced Ca^2+^ responses. Together, these results suggest that PA-MPs increase cellular stiffness and reduce mechanically evoked Ca^2+^ signaling via PIEZO1.

**Fig. 2.**
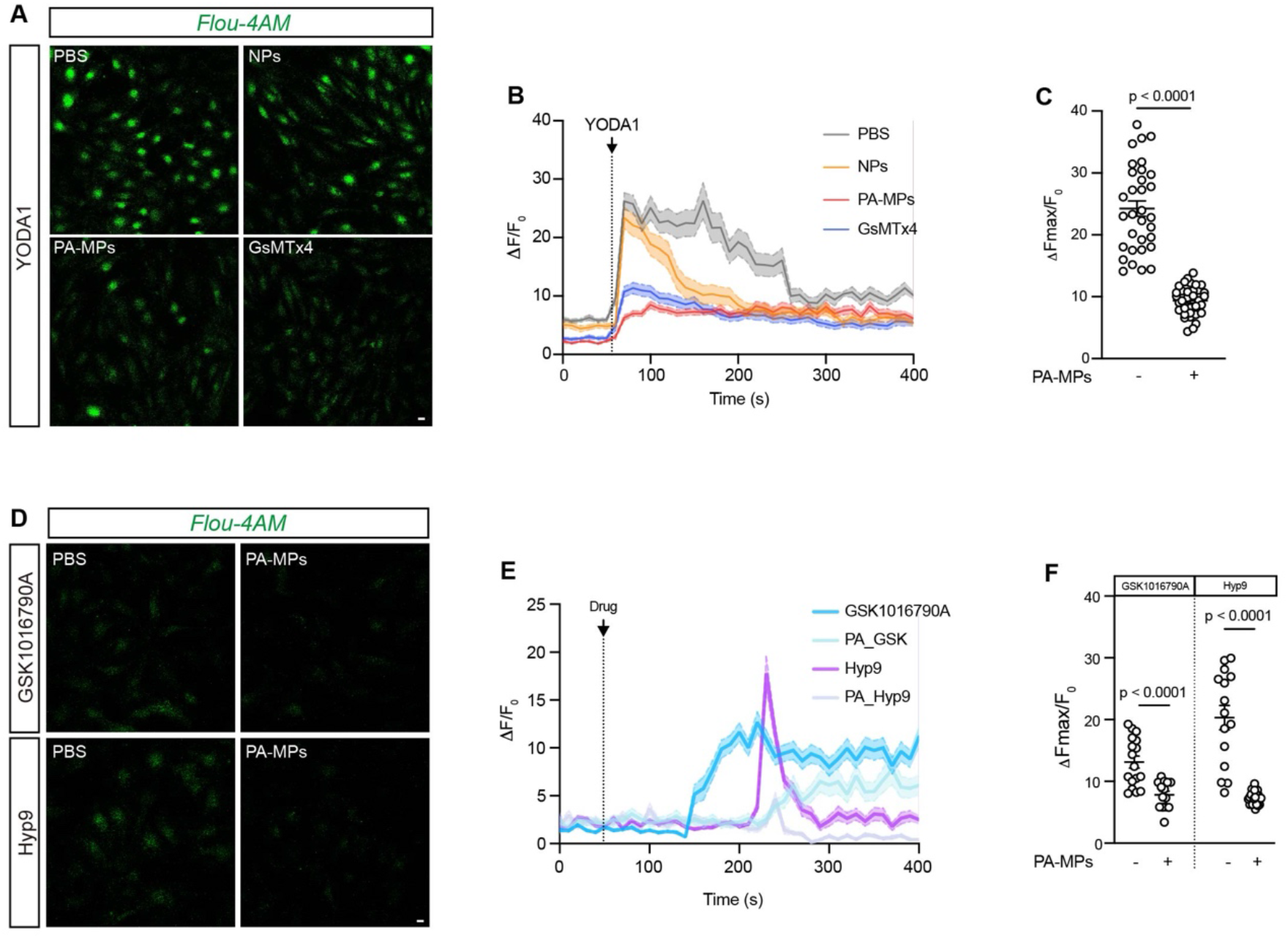
PA-MPs selectively impair Piezo1-mediated Ca^2+^ signaling and are not compensated by TRPV4 or TRPC6 activation. (**A**) Live-cell calcium imaging of Fluo-4 AM-loaded HAECs, revealing intracellular Ca^2+^ responses to the Piezo1 agonist Yoda1. Robust Ca^2+^ influx was observed in control and NPs-treated cells, whereas PA-MPs and GsMTx4-treated cells exhibited markedly attenuated Yoda1-induced Ca^2+^ responses. (**B**) Representative time-lapse images and corresponding fluorescence traces showing intracellular Ca^2+^ influx before and after Yoda1 application. Time-course plots illustrate normalized fluorescence intensity (ΔF/F_0_) over the recording period. (**C**) Quantification of maximal Ca^2+^ response following Yoda1 treatment, expressed as ΔFmax/F_0_, in individual ECs (n = 32-34 cells). (**D, E**) Pharmacological activation of mechanosensitive ion channels TRPV4 and TRPC6 using their selective agonists, GSK1016790A and Hyp9, respectively (**D**), failed to restore Ca^2+^ signaling in PA-MPs treated cells (**E**). (**F**) Quantification of maximal Ca^2+^ responses (ΔFmax/F_0_) in individual ECs following stimulation with GSK1016790A or Hyp9 (n = 14-24 cells).

### PA-MPs modulate mechanotransduction- and Notch-associated junctional dysfunction in endothelial cells

The transcriptional profile underlying PA-MPs-induced EC dysfunction was assessed by bulk RNA sequencing following exposure of HAECs to NPs or PA-MPs (Fig. 3 A to D). PA-MPs downregulated genes governing endothelial mechanotransduction (*ITGB1, ITGB3, ATP2B1, ATP2B2, CAV1, YAP1*), barrier integrity (*ROCK1/2, TJP1, F11R, ANGPT2*), and Hippo–Notch signaling (*RBPJ, MAP4K3, LATS1/2, MOB1A/B, WWTR1, ADAM10*), while it upregulated genes with inflammatory and innate immune pathways associated cytokines and chemokines (*IL6, IL1B, CCL2, CXCL10, ICAM1*), viral RNA-sensing pathways (*TLR3, RIGI, IFIH1*), IRF-dependent interferon signaling and a broad interferon-stimulated associated genes (*MX, OAS, ISG15, RSAD2)*, and *IFIT* family members (Fig. 3D). Gene ontology (GO) analysis corroborated these findings, revealing coordinated repression of transcriptional regulation, protein modification, and autophagy pathways, alongside robust activation of antiviral and interferon responses (Fig. 3, B and C). Then, we verified the transcriptomic profiling that PA-MPs reduced NOTCH pathway-associated genes (*DLL4*) and endothelial cell junctional genes (*ZO1* and *CLDN5*), and increased NOTCH-related genes (*HES1* and *HEY1*) and an inflammatory marker (*TNF*) by quantitative PCR (fig. S2). However, *Piezo1* gene expression was not similar between control and PA-MPs. Collectively, these transcriptomic data indicate that PA-MPs exposure enhances pro-inflammatory and interferon-related gene expression and suppresses pathways associated with mechanotransduction and endothelial junctional integrity. These transcriptomics phenotypes suggest that PA-MPs induced endothelial barrier dysfunction with activation of sterile inflammatory signaling. Consistent with the transcriptomic profile, PA-MPs exposure reduced protein expression levels of the Notch intracellular domain (NICD), a key mediator of Piezo1-dependent regulation of endothelial junctional integrity (*18, 27*), TRPV4, and TRPC6, whereas PIEZO1 protein expression remained unchanged (Fig. 4, A and B). The preceding experiments demonstrated that the Peizo1 agonist Yoda1 enhances Piezo1 function in control conditions but fails to restore signaling in PA-MPs-treated cells. This observation prompted us to examine whether Piezo1 activation also influences downstream signaling at the level of protein abundance. Consistent with this possibility, Yoda1 treatment increased NICD protein levels in PBS and NP conditions, whereas this effect was strikingly attenuated in the presence of PA-MPs (Fig. 4, C and D).

**Fig. 3.**
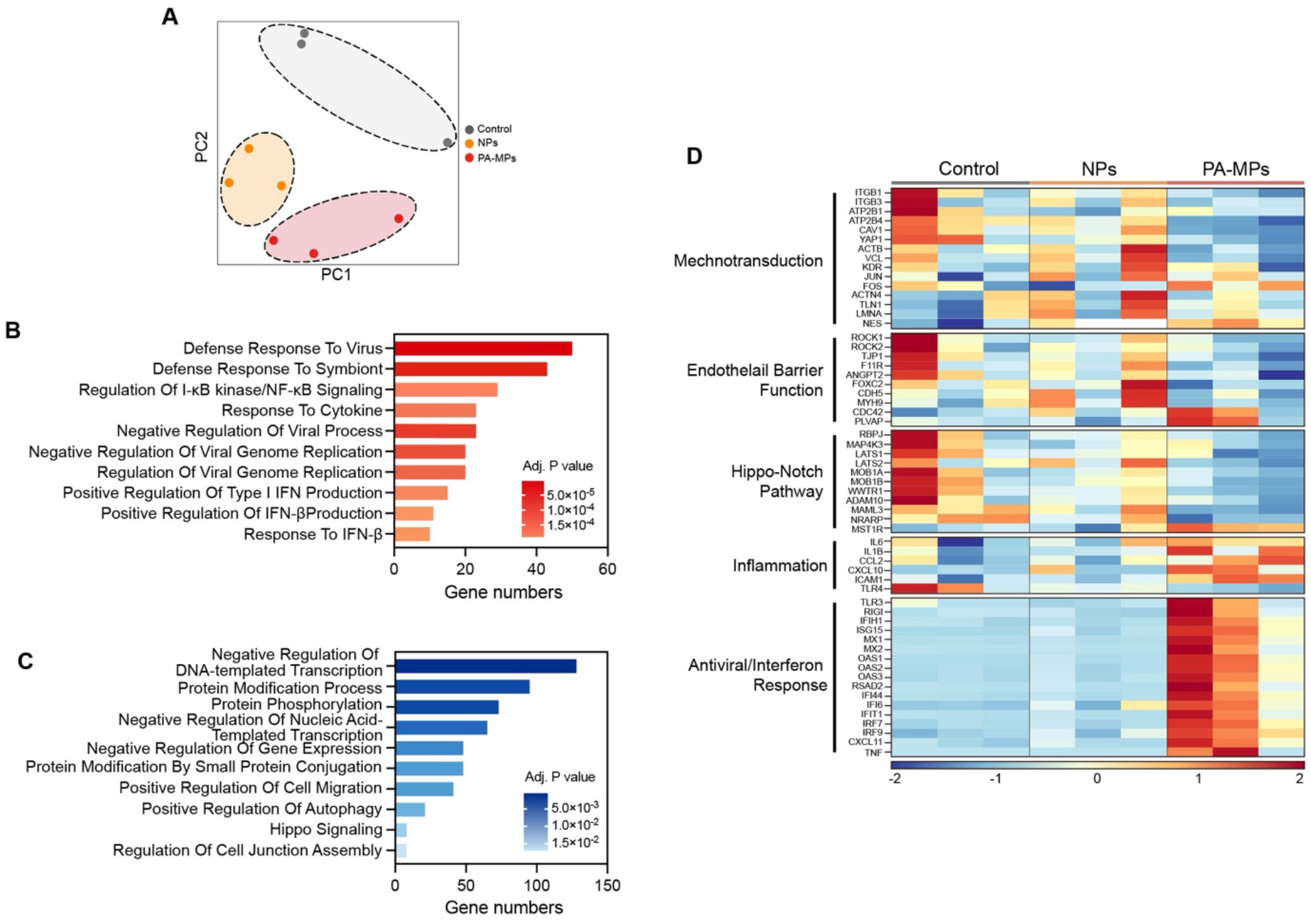
PA-MPs induce transcriptional reprogramming in human aortic endothelial cells (HAECs). (**A**) Principal component analysis (PCA) showing distinct clustering of control, NPs, and PA-MPs-treated endothelial cells based on global transcriptomic profiles. (**B, C**) Gene ontology (GO) enrichment analysis of differentially expressed genes (DEGs) reveals significantly enriched biological processes that are upregulated (**B**) or downregulated (**C**) following PA-MPs exposure. (**D**) Heatmap of DEGs in HAECs treated with PBS, NPs, or PA-MPs, grouped by functional categories (n = 3 independent biological replicates per condition).

**Fig. 4.**
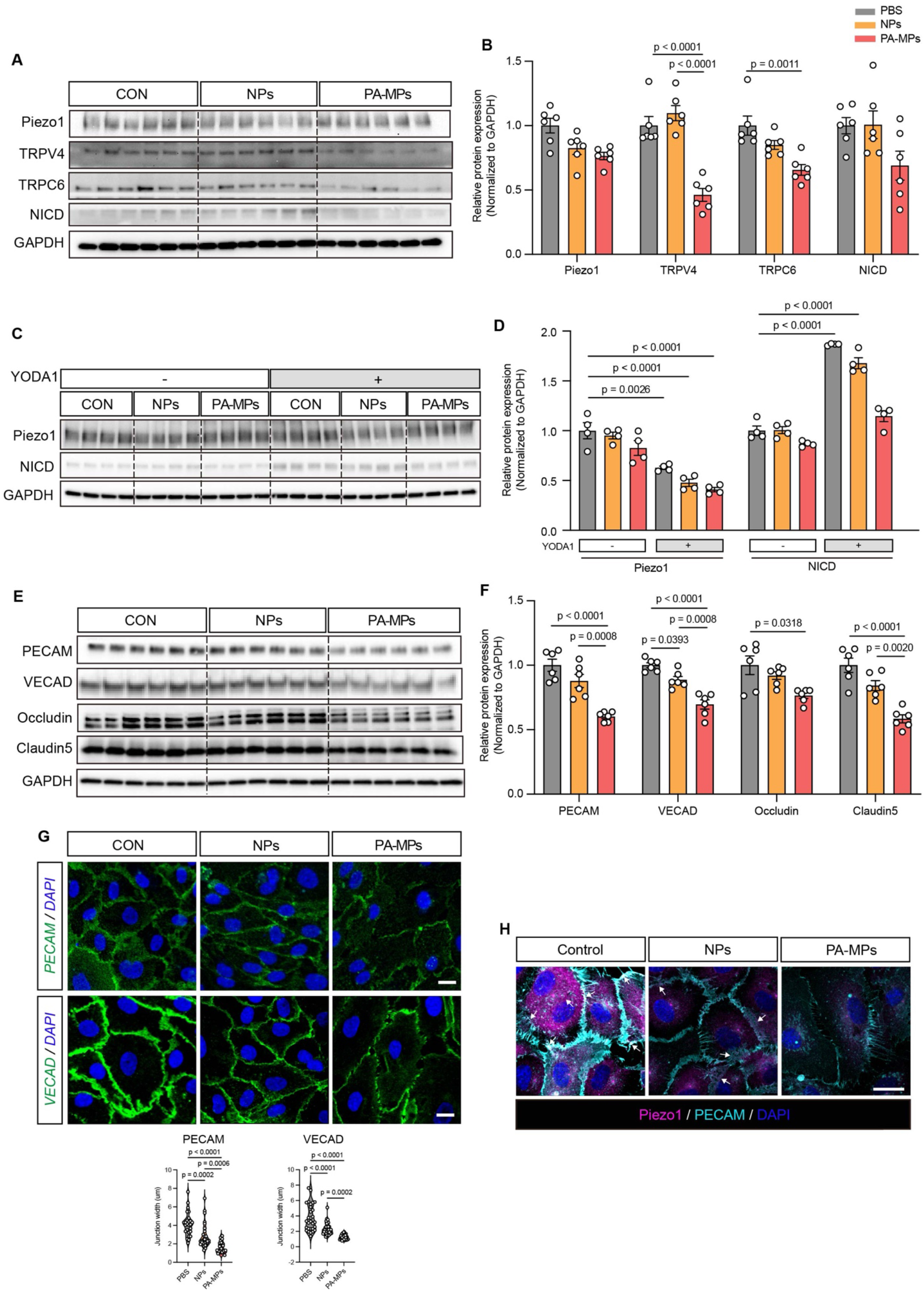
PA-MPs exposure functionally uncouples mechanosensitive ion channels from Notch signaling and barrier maintenance. (**A**) Representative immunoblots illustrating expression of mechanosensitive ion channels (Piezo1, TRPV4, and TRPC6) and the active Notch signaling component Notch intracellular domain (NICD) in HAECs. (**B**) Quantitative analysis revealed that PIEZO1 protein abundance remained unchanged across all experimental conditions, whereas TRPV4 and TRPC6 expressions were significantly reduced following PA-MPs exposure. Consistent with impaired mechanotransduction, NICD levels were markedly decreased in PA-MPs treated cells, implicating suppression of Notch signaling. (**C** and **D**) Pharmacologic activation of Piezo1 with Yoda1 increased NICD levels in PBS and NPs-treated cells but failed to restore NICD expression in PA-MPs exposed cells This occurred despite preserved PIEZO1 protein abundance, indicating a functional uncoupling of Piezo1 activation from downstream Notch signaling. (**E** and **F**) Immunoblot analyses of adherens junction proteins (PECAM-1 and VE-cadherin) and tight junction proteins (Occludin and Claudin-5) demonstrated substantial downregulation in PA-MPs–exposed cells compared with PBS and NPs, consistent with compromised endothelial barrier integrity. (**G**) Immunofluorescence imaging revealed junctional disorganization and narrowing, evidenced by the disruption of the continuous staining of PECAM-1 and VE-cadherin at cell–cell borders. (**H**) HAECs were stained for Piezo1(magenta) and PECAM1 (cyan) following treatment under the indicated conditions. White arrows highlight regions of Peizo1-PECAM1 colocalization at the plasma membrane. Scale bar: 20 µm.

PA-MPs reduced the expression of adherent junctional proteins (PECAM-1 and VE-cadherin) and tight junction proteins (Occludin and Claudin-5) (Fig. 4, E to G), and increased endothelial permeability as measured by transwell assay (fig. S3). In addition, substantial colocalization of Piezo1 with platelet endothelial cell adhesion molecule (PECAM)-1 at the endothelial plasma membrane was observed under control conditions, which was progressively diminished following NPs and PA-MPs exposure (Fig. 4H), suggesting that microplastics compromise EC junctional protein integrity and increase cell-cell permeability. Taken together, PA-MPs impair Piezo1-dependent mechanotransduction, attenuate Notch pathway activation, and weaken endothelial junctional integrity.

### PA-MPs inhibit Piezo1-dependent Notch signaling *in vivo*

To extend our mechanistic findings to an *in vivo* context, we utilized the zebrafish (*Danio rerio*) model to investigate how the disruption of endothelial signaling triggers vascular barrier compromise and facilitates the subsequent entry of particles into the circulation. Zebrafish offer a unique platform that enables high-resolution visualization of vascular barrier integrity and spatiotemporal tracking of circulating particles under physiologically relevant exposure conditions, providing an ideal system to assess the systemic consequences of barrier function (*28, 29*).

First, we test the ability of PA-MPs to translocate and disrupt the intestinal-vascular barrier, PA-MPs were administered via micro-gavage to *Tg(flk1:EGFP)* zebrafish larvae at 5 days post fertilization (dpf), enabling direct visualization of endothelial integrity (Fig. 5A and fig. S4, A and B). After micro-gavage, the survival rate was monitored for 4 days post-gavage (dpg). PA-MPs led to a progressive decline in survival, reaching 67% by day 4, compared with survival rates of 98% and 92% in the PBS- and NPs-treated groups, respectively (fig. S4C). Endothelial barrier function was evaluated at 6 or 24 hours of exposure to PA-MPs, revealing robust disruption of the vascular barrier within the caudal vein plexus (CVP). At 24 h post-exposure, vascular barrier compromise was detected in 0% of PBS controls, 25% of NPs-treated, and 75% of PA-MPs treated larvae (Fig. 5, B and C). Supplementary movie 2 shows that PA-MPs circulate within the bloodstream of zebrafish. In this context, PA-MPs promotes to greater intestinal-vascular barrier dysfunction leading to penetration of PA-MPs into the circulating system in zebrafish.

**Fig. 5.**
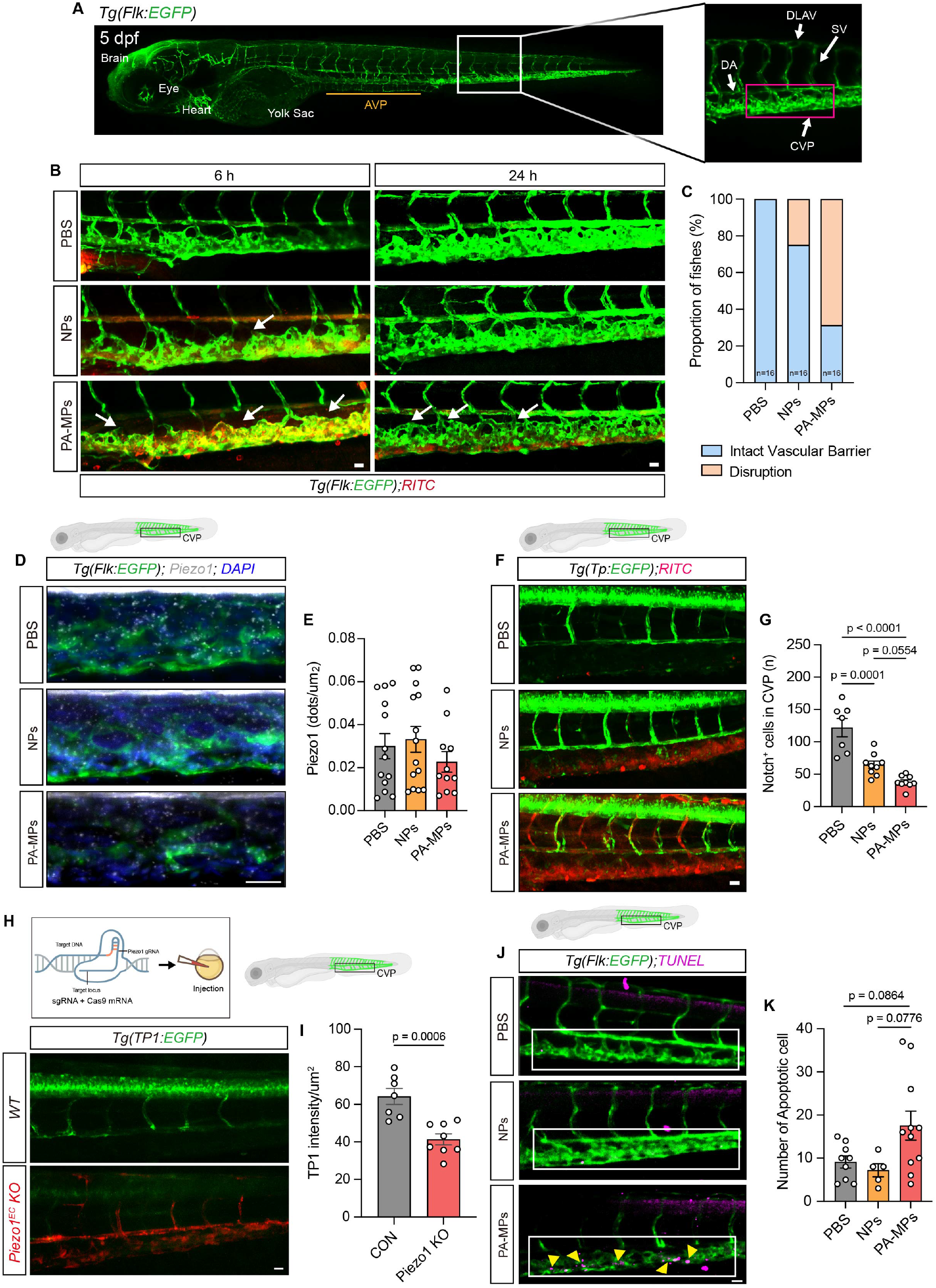
PA-MPs disrupt Piezo1–Notch signaling, compromise vascular barrier integrity, and promote endothelial apoptosis *in vivo*. (**A**) Schematic and representative confocal images of a transgenic *Tg(Flk1:EGFP)* zebrafish embryo at 5 days post-fertilization (dpf), highlighting the vascular anatomy. A yellow line delineates the anterior venous plexus (AVP). White arrows indicate the dorsal aorta (DA), caudal venous plexus (CVP), posterior caudal vein (PCV), segmental vessels (SVs), and dorsal longitudinal anastomotic vessel (DLAV). The PCV region analyzed is demarcated by a magenta box. (**B**) *Tg(Flk1:EGFP)* larvae were subjected to micro-gavage with PBS, NPs, or PA-MPs and imaged at 6 or 24 hours post-gavage (hpg). PA-MPs exposure induced marked vascular barrier disruption within the CVP, evidenced by endoluminal retention of RITC fluorescence and structural abnormalities. White arrows denote regions of compromised endothelial integrity. (**C**) Quantification of larvae exhibiting endoluminal RITC fluorescence and vascular morphological defects demonstrates a significantly higher incidence of barrier disruption following PA-MP exposure (n = 16 per condition). (**D**) Representative confocal images of *Tg(Flk1:EGFP)* zebrafish micro-gavaged at 5 dpf and analyzed at 6 dpf. Fluorescent in situ hybridization (FISH) for Piezo1 mRNA within the CVP revealed no detectable differences in transcript abundance across PBS, NPs, and PA-MPs treatment groups. (**E**) Quantification of *Piezo1* transcript density, expressed as Piezo1-positive puncta per region of interest (ROI; µm^2^), confirmed preserved transcriptional expression (n = 10 per group). (**F**) Representative images of *Tg(Tp1:EGFP)* zebrafish, in which EGFP reports transcriptional activity of the Notch-responsive Tp1 promoter. PA-MPs exposure markedly reduced EGFP signal within the CVP, indicating suppressed Notch1 signaling. (**G**) Quantification of Tp1^+^ (Notch1-active) endothelial cells in the CVP demonstrates a significant reduction following PA-MP exposure (n = 7-9 per group). (**H** and **I**) Functional validation of Piezo1–Notch coupling. Endothelial-specific Piezo1 knockout zebrafish generated via CRISPR-Cas9 and crossed with the *Tg(Tp1:EGFP)* reporter line exhibited a pronounced decrease in Notch activity relative to control embryos. (**J** and **K**) TUNEL staining revealed significantly increased endothelial apoptosis in the CVP of PA-MP-treated larvae compared with PBS and NPs controls (n = 5-9 per group). Yellow arrowheads indicate apoptotic cells. Scale bars: 20 µm unless otherwise indicated.

Subsequently, we examined the effect of PA-MPs on *piezo1* transcription *in vivo* by quantifying Piezo1 RNA expression in gut vascular endothelial cells. *In situ* hybridization in *Tg(flk1:EGFP)* larvae demonstrated no significant differences in *piezo1* mRNA levels in CVP following exposure to NPs or PA-MPs compared to PBS controls (Fig. 5, D and E), which is consistent with the maintained expression of PIEZO1 mRNA (fig. S2) and protein (Fig. 4A). Notch signaling activity, evaluated by using *Tg(Tp1:EGFP)* reporter line for the Notch-responsive (Tp1^+^) endothelial cells, was diminished in the CVP following PA-MPs (Fig. 5, F and G). To evaluate the EC Piezo1 in gut vascular permeability and Notch signaling, EC-specific *piezo1*-deficient *Tg(Tp1:EGFP)* zebrafish embryos were generated via a CRISPR-Cas9-based strategy (Fig. 5H) to investigate the role of Piezo1 and Notch signaling *in vivo*. Developmental progression, survival, and gene-editing efficiency following EC-specific *piezo1* loss-of-function were evaluated in zebrafish embryos (fig. S5, A to D). *Piezo1* deficiency resulted in a significant reduction in Tp1-driven EGFP fluorescence compared with scrambled control groups (Fig. 5I), phenocopying the suppression of Notch signaling induced by PA-MPs exposure. In contrast, pharmacological blockade of Notch signaling with the γ-secretase inhibitor DAPT did not affect *piezo1* transcript levels in the CVP (fig. S6, A to C), consistent with a unidirectional regulatory relationship in which Piezo1-mediated calcium influx regulates downstream Notch activation. Also, TUNEL staining showed elevated apoptosis within the CVP of PA-MPs-exposed larvae relative to PBS and NPs-treated groups (Fig. 5, J and K).

### Photoaged microplastics disrupt Piezo1-dependent calcium signaling in neural and cardiac tissues and impair locomotor performance

While previous studies have described functional outcomes following systemic administration of MPs, typically achieved through direct intravenous or intratracheal delivery in rodent models, such approaches circumvent physiological barrier interfaces relevant to environmental exposure (*30, 31*). This methodological gap leaves the dynamics of barrier crossing and subsequent systemic distribution insufficiently defined. Here, we demonstrate that PA-MPs breach the gut-vascular barrier in zebrafish, directly linking environmentally relevant exposure pathways to systemic functional impairment. Given this translocation into the circulation, we subsequently examined functional perturbations in distal organ systems. With this rationale, genetically encoded calcium indicators with high sensitivity and cell-type specificity were used to monitor pan-neuronal activity in *Tg(elavl3:GCaMP6s)* larvae (*32*) (Fig. 6A and movie S3) and cardiomyocyte calcium dynamics in *Tg(myl7:gCaMP4*.*1*^*LA2124*^) larvae (*33*) (Fig. 6C and movie S4). In Fig. 6B, ΔF/F_0_ traces depict calcium transient activity associated with neuronal firing events. PA-MPs exposure substantially attenuated the amplitude and frequency of neuronal calcium spikes. Comparable effects were observed in the heart, where calcium transients in 6 dpf *Tg(myl7:gCaMP4*.*1*^*LA2124*^) larvae were reduced following PA-MPs treatment (Fig. 6, C and D). As shown in fig. S7, Yoda1 increased calcium transient amplitude by approximately 30% in PBS and NPs treated groups, whereas no recovery was detected in PA-MPs-exposed larvae (fig. S7 and movie S5). Thus, these data indicate that PA-MPs exposure disrupts both basal and evoked calcium dynamics in neuronal and cardiac tissues *in vivo*. PA-MPs translocation beyond the vascular barrier coincided with reduced calcium signaling in these tissues, consistent with particle presence in distant organs and disruption of calcium-dependent physiological processes.

**Fig. 6.**
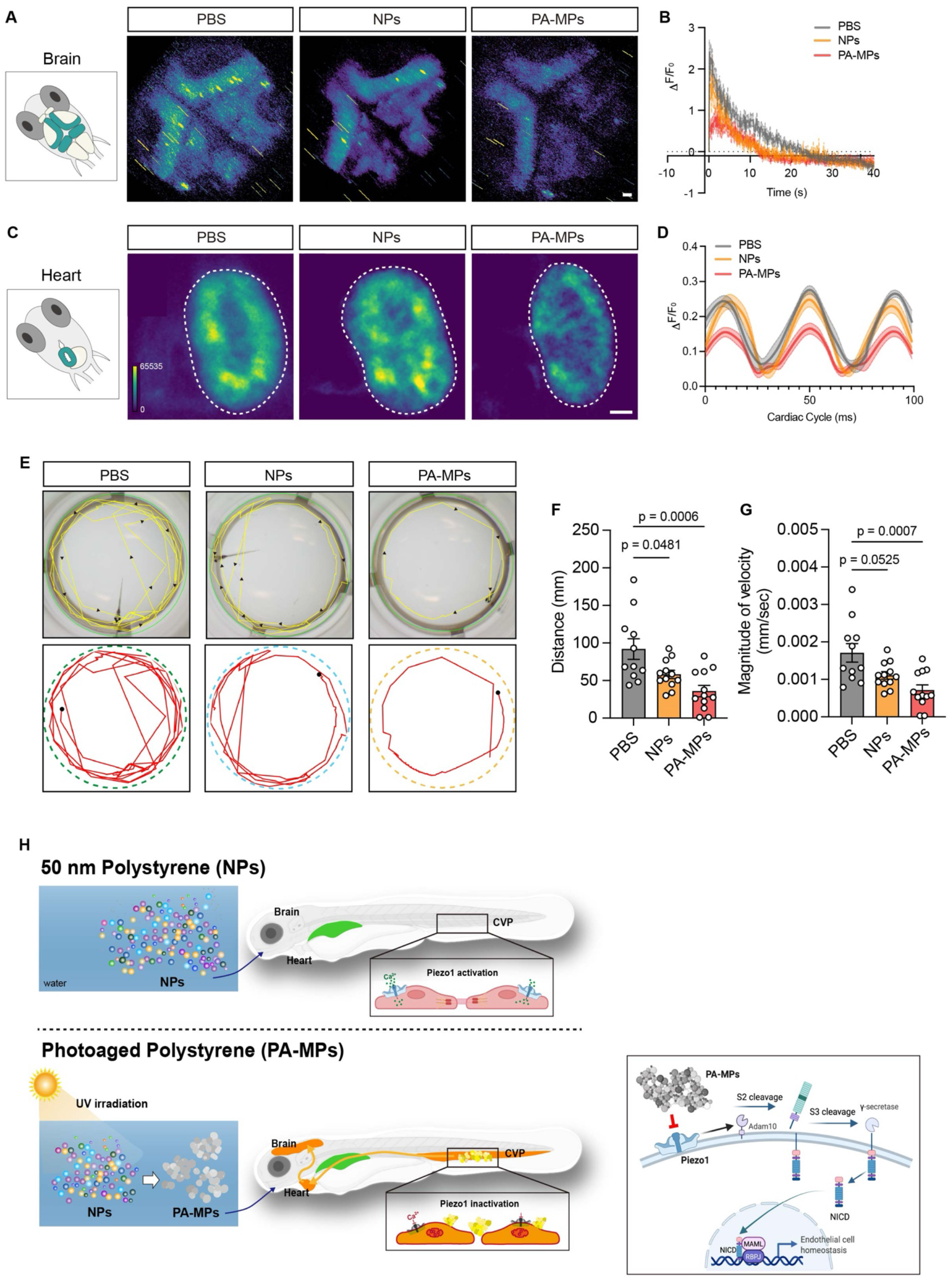
PA-MPs attenuate calcium influx in the brain and heart, impairing zebrafish locomotive function. (**A**) Representative calcium imaging of neuronal activity in *Tg(elavl3:GCaMP6s)* zebrafish larvae at 6 dpf following micro-gavage with PBS, NPs, or PA-MPs. (**B**) Quantitative analysis of calcium transients within the optic tectum, with GCaMP fluorescence signals integrated across the entire brain region of interest. PA-MPs exposure significantly suppressed neuronal calcium activity compared with PBS and NP controls (n = 9-15 per group). (**C**) Representative radiometric calcium imaging of cardiac activity in *Tg(myl7:gCaMP4*.*1*^*LA2124*^) zebrafish larvae at 6 dpf following PBS, NPs, or PA-MPs exposure. (**D**) Quantified myocardial calcium transient traces demonstrating attenuated Ca^2+^ dynamics in PA-MP-treated larvae (n = 4-7 per group). (**E**) Representative cumulative swimming trajectories recorded during a 5-minute locomotor assay, illustrating impaired swimming performance in PA-MP-exposed larvae relative to controls. (**F** and **G**) Quantification of total swimming distance (**F**) and mean swimming velocity (**G**) confirms significant locomotor deficits following PA-MPs exposure (n = 12 per group). Scale bar: 20 µm. (**H**) Model of PA-MP-induced endothelial Piezo1-Notch dysfunction. Photoaged particles impair Piezo1-dependent Ca^2+^ entry without altering expression, leading to reduced Notch signaling, barrier disruption, systemic dissemination, and associated functional alterations in zebrafish.

Finally, the potential behavioral consequences of disrupted neuronal and cardiac calcium signaling were evaluated using locomotor activity as a sensitive and early indicator of neurotoxicity and ecological relevance (*34*). Interestingly, larvae exposed to PA-MPs exhibited significantly reduced swimming distance and velocity compared with PBS-treated controls, whereas intact MPs had a lesser impact on locomotive behavior (Fig. 6, E to G). Behavioral shifts emerged in parallel with changes in neuronal and cardiac calcium signaling, indicating a possible relationship between disrupted calcium dynamics and behavioral performance.

## Discussion

Microplastics and nanoplastics (MNPs) are recognized as pervasive environmental contaminants, arising from diverse sources and undergoing continuous physicochemical transformation under natural conditions (*35, 36*). The environmental aging process, particularly ultraviolet-induced photooxidation, modifies particle size, surface chemistry, hydrophilicity, and aggregation behavior, consequently modulating their biological reactivity and toxicity profiles (*37-39*). These transformations enhance oxidative potential, alter colloidal stability, and increase interactions with biological interfaces, ultimately amplifying adverse effects in aquatic and biological systems (*4, 22, 40*).

Our SEM analyses show that prolonged UV exposure induces pronounced fragmentation and self-aggregation of polystyrene nanoplastics. These structural changes, aggregation, enlargement, and roughness of surface, were more dramatically observed in E3 medium than in DDW, which underscores how the surrounding aqueous composition strongly influences photooxidative transformation and particle reorganization. To our knowledge, these findings provide the first direct evidence that environmentally relevant conditions exacerbate photooxidation-associated structural remodeling of nanoplastics compared with simplified aqueous systems.

Although it has been well established that aggregation, transport, and toxicological properties of MNPs, most studies have relied on pristine or commercially manufactured particles that incompletely represent environmentally aged materials (*2*). Also, there have been detected in the brain (*13, 30*), heart (*31*), liver (*41*), and vascular tissues (*42*) across aquatic models and mammalian systems, as well as their presence in human cardiovascular tissues (*12*), where particle detection correlates with elevated risks of myocardial events. However, the mechanisms by which PA-MPs exposure promotes systemic translocation remains unresolved. Thus, in this study, we focused on Piezo1 signaling and found that exposure to PA-MPs attenuated Piezo1-dependent Ca^2+^ entry and downstream Notch1 activation in both zebrafish endothelial cells and HAECs, despite preservation of PIEZO1 transcript and protein and levels. These findings therefore indicate functional uncoupling of mechanosensitive signaling rather than transcriptional or translational suppression. Accordingly, partial restoration of NICD levels by Piezo1 agonist treatment further implicates Piezo1-mediated Ca^2+^influx as an upstream regulator of Notch-dependent signaling required for endothelial barrier homeostasis (*18, 27*). Transcriptomic profiling further revealed a coordinated repression of pathways that regulate mechanotransduction, cytoskeletal organization, and junctional maintenance, paired with the induction of inflammatory, antiviral, and interferon-responsive programs; this shift characterizes a transition from mechano-adaptive endothelial homeostasis toward a stress-dominated, pro-apoptotic state (*43-45*).

We also establish that PA-MPs, but not pristine NPs, compromise endothelial integrity at the gut-vascular barrier, enabling their translocation into the systemic circulation and subsequent accumulation in distant organs with detrimental functional consequences. Specifically, in the zebrafish model, PA-MPs disrupted both the intestinal epithelial barrier and the underlying vascular endothelium, which promotes observable gut-vascular leakage and widespread systemic translocation of the particles (*46, 47*). In this context, PA-MPs exposure disrupted neuronal and cardiomyocyte calcium dynamics and impaired locomotor performance. These observations are consistent with recent reports that microplastics accumulate across multiple organs in experimental models (*30, 31, 41, 42*) and humans (*12, 13*). Such widespread accumulation necessitates translocation via the bloodstream, supporting the concept that endothelial barrier dysfunction may present a critical gateway for systemic exposure. We propose that the impairment of endothelial mechanosensitive ion channel signaling, with Piezo1 as a key regulator. Previous studies have shown that PECAM-1 interacts with Piezo1 to direct its localization at endothelial cell-cell junctions (*48*), and that Piezo1 function has been predominantly examined in the context of mechanical forces or altered hemodynamic flow (*49-51*). Our findings extend current mechanistic understanding and show that MPs, as an environmental stressor, microplastics, impair mechanosensitive ion channel function in endothelial cells. The functional decoupling of Piezo1 from its junctional signaling context, caused by this impairment, resulted in weakened vascular barrier integrity.

Several limitations of this study should be acknowledged. First, the experimental exposure conditions employed here may exceed estimated human environmental burdens, and direct evidence for MNPs penetration into human vascular tissues remains limited. Second, although zebrafish models provide unique advantages for *in vivo* imaging and functional interrogation (*52*) of vascular and systemic processes, they may not fully recapitulate the complexity of human exposure scenarios, such as chronic low-dose exposure, heterogeneous plastic compositions, and the influence of pre-existing cardiometabolic risk factors. Overcoming these limitations will require future studies employing environmentally relevant plastic mixtures, prolonged low-level exposures, and complementary mammalian or human-based systems, including organoid models, *ex vivo* vascular tissues, and population-based approaches. Further investigations will be critical to ascertain whether the disruption of endothelial mechanotransduction observed here represents a conserved pathophysiological response across diverse species and varying environmental exposure contexts.

Despite these limitations, our findings highlight an underappreciated dimension of plastic toxicity, demonstrating that environmental aging processes can fundamentally transform particle bioactivity by specifically impairing endothelial mechanosensitive signaling. These insights suggest that current risk assessment approaches must adapt to incorporate aging-related endpoints and underscore the importance of global plastic mitigation to protect both vascular homeostasis and public health.

## Methods

### Preparation of photoaged-polystyrene microplastics (PA-MPs)

Polystyrene nanoplastics (50 nm) were obtained from Degradex by Photophorex (Hopkinton, MA). Upon receipt, particles were either stored in their original aqueous suspension containing 0.1% Tween-20 or subjected to ultraviolet (UV) photoaging. Nanoplastics were pipetted onto sterile glass dishes submerged in E3 medium for photoaging and placed in a biosafety cabinet with a UV lamp. Based on established protocols, continuous UV exposure was applied for 2 or 4 weeks to simulate environmental aging (*53*). Throughout the aging process, water was periodically replenished to maintain hydration. Non-aged (fresh) particles were stored at 4°C in the dark and shielded from UV light until use in experimental treatments.

### Scanning Electron Microscope (SEM)

SEM imaging was performed following a procedure similar to a previously reported method using a Nova Nano 230 field-emission scanning electron microscope (FEI) (*54*). Particles obtained from the preceding experimental step were drop-cast onto a silicon wafer and air-dried. The wafer was then briefly dipped in water, dried again, and sputter-coated with an approximately 3 nm gold layer using a Pelco SC-7 sputter coater (40 s) before imaging.

### Atomic force microscopy (AFM)

AFM measurements were performed following a previous described protocol (*55*). Human aortic endothelial cells (HAECs; Lonza Inc.) were cultured in vitro to approximately 60% confluence with or without PA-MPs treatment and analyzed using a JPK NanoWizard 4a BioScience AFM. Indentation was performed with a Bruker SAA-SPH-1UM probe (spring constant *k* ≈ 0.25 N m^−1^; the exact *k* of each probe was calibrated by laser Doppler velocimetry). Force–distance curves were recorded, and Young’s modulus was determined by fitting the indentation data to the Hertz/Sneddon model using JPK Data Processing software (v 3.4).

### Cell culture and Western blotting

HAECs were cultured in endothelial basal medium-2 supplemented with EGM-2 SingleQuots (Lonza Inc.) at 37°C in a 5% CO_2_ atmosphere. Cells were passaged at 70-80% confluence, and passages 4-6 were used for experiments (*56, 57*). HAECs were treated with 9 ug/ml of NPs or PA-MPs for 24 h. Following treatment, cells were lysed in RIPA buffer supplemented with protease/phosphatase inhibitors and EDTA.

Protein lysates (30 µg per sample) were mixed with 6x sample buffer, resolved on 4-20% SDS-PAGE gels (Bio-Rad, #5641094), and transferred to PVDF membranes using a semi-dry transfer system (Trans-Blot Turbo, Bio-Rad). Membranes were blocked with 5% skim milk for 1 h at room temperature and incubated overnight at 4°C with primary antibodies against Piezo1, PECAM, VECAD, NICD1, Claudin5, 4-HNE, and GAPDH (table S1). After incubation with HRP-conjugated secondary antibodies (goat anti-rabbit, goat anti-mouse, and rabbit anti-goat), protein bands were visualized by chemiluminescence and quantified using ImageJ (NIH).

### Measurement of Intracellular Ca^2+^ concentration

Endothelial cells (ECs) were seeded on laminin-coated 35 mm glass-bottom dishes and exposed to NPs or PA-MPs for 24 h. Cells were then incubated with 2 µM Fluo-4 AM and 0.024% Pluronic F-127 (ThermoFisher) in HBSS solution for 30 min at 37°C. After washing, dishes were transferred to a temperature-controlled chamber (37°C) on a confocal microscope (Leica SP8, 488 nm excitation). Fluorescence signals were recorded from ∼30 cells per dish, with 3 biological replicates. Intracellular Ca^2+^ variations were expressed as F/F_0_, where F represents fluorescence during recording and F_0_ represents the baseline fluorescence. To assess Piezo1 activation, 2 uM Yoda1 (Sigma-Aldrich, MO, USA) was applied after a 2 min baseline recording (*58*). For ion channel blockade, cells were pretreated with 5 uM GsMTx4 for 30 min prior to Yoda1 stimulation. To examine additional mechanosensitive transducers, TRPV4 and TRPC6 were activated using their respective agonists, GSK1016790A (50 nM) and Hyp9 (10 uM).

### RNA-seq analysis

For bulk RNA-seq analysis, HAECs were treated with PBS, NPs, or PA-MPs in three independent biological replicates per condition. Total RNA was isolated using the Qiagen RNeasy Mini Kit according to the manufacturer’s instructions. RNA integrity and quality were assessed using the RNA High Sensitivity Assay on a TapeStation 2200 (Agilent Technologies). RNA-seq libraries were prepared using the KAPA mRNA HyperPrep Kit following the manufacturer’s protocol. Briefly, the workflow included poly(A)+ mRNA enrichment and fragmentation, followed by first-strand cDNA synthesis using random priming and second-strand synthesis incorporating dUTP to generate double-stranded cDNA. Subsequent steps included end repair, A-tailing, adaptor ligation, and PCR amplification, with unique adaptors used for multiplexing samples within a sequencing lane. Libraries were diluted, pooled, and sequenced on an Illumina NovaSeq X Plus platform with a paired-end 2 x 50 bp configuration. Sequencing quality control was performed using Illumina Sequencing Analysis Viewer (SAV), and base calling and demultiplexing were carried out using Illumina BCL Convert v4.2.7. Raw paired-end FASTQ files generated by the UCLA sequencing core were subjected to quality control and reads shorter than 20 base pairs or with an average Phred quality score below 20 were excluded. High-quality reads were aligned to the human reference genome (GRCh38.p14, release 47) using the STAR RNA-seq aligner (*59*). Gene-level read counts were quantified with featureCounts to generate the expression matrix. Raw count data were log1p-normalized prior to dimensionality reduction, and Principal Component Analysis (PCA) was performed and visualized using Scanpy. Differential gene expression analysis was conducted on raw count matrices using PyDESeq2. Functional enrichment of differentially expressed genes was assessed by Gene Set Enrichment Analysis using GSEpy via the EnrichR application programming interface. For visualization, gene-wise Z-scores were calculated and used to generate heat maps with Scanpy’s plotting utilities (*60, 61*).

### Quantitative PCR (qPCR)

Total RNA was isolated from NPs and PA-NPs-treated cells using the RNeasy Mini Kit (Qiagen, #74104) according to the manufacturer’s protocol (*62*). RNA concentration and purity were assessed with a NanoDrop spectrophotometer (ThermoFisher, #ND-ONE-W). cDNA was synthesized from 1 μg RNA using the iScript Reverse Transcription Supermix (Bio-Rad). qPCR was performed using SYBR Green Supermix (Bio-Rad) on a CFX96 Touch Real-Time PCR Detection System (Bio-Rad). Primer sequences are provided in the Table. Thermal cycling conditions were as follows: initial denaturation at 95°C for 30 sec, followed by 40 cycles of 95°C for 15 and 60°C for 30 sec. Gene expression was normalized to GAPDH as an internal control, and relative expression was calculated using the 2^-ΔΔCt^ method. Each reaction was run in duplicate with at least two biological replicates. Primer sequences used for qPCR analysis are listed in table S2.

### Immunofluorescence

Cells were fixed with 4% paraformaldehyde (PFA; Sigma-Aldrich, MO, USA) for 10 min at room temperature and stored at -80°C until use. For staining, cells were thawed, permeabilized with 0.1% Triton X-100, and blocked with 5% donkey serum for 1 h at room temperature. Samples were then incubated with primary antibodies against PECAM or VECAD, followed by 594 Alexa Fluor-conjugated secondary antibodies (ThermoFisher) for 1 h at room temperature. Nuclei were counterstained with DAPI, and coverslips were mounted using ProLong Diamond Antifade Mountant (ThermoFisher). Images were acquired on a confocal microscope (Leica SP8, Wetzlar, Germany) using a 20x objective. All images were collected with identical acquisition settings and corrected for background fluorescence.

### Zebrafish husbandry

All zebrafish experiments were conducted under protocols approved by the UCLA Institutional Animal Care and Use Committee (IACUC). Under standard conditions, Zebrafish were maintained at 28.5°C in the UCLA Zebrafish Core Facility. Embryos were raised in E3 medium containing 0.05% methylene blue (Sigma-Aldrich, MO, USA) to prevent fungal growth and 0.003% phenylthiourea (PTU; Sigma-Aldrich, MO, USA) to inhibit melanogenesis (*63*).

The AB strain was used as the wild type. The following transgenic lines were used: *Tg(Flk: EGFP*) (*64*), *Tg*(*Tp1:EGFP*) (*29*), *Tg*(*elavl3:GCaMP6s*) (*32*), and *Tg*(*myl7:gCaMP4*.*1*^*LA2124*^) (*33*). To assess Piezo1-Notch pathway interactions, larvae from the *Tg(Tp1:EGFP)* line were treated with 50 μM DAPT, a γ-secretase inhibitor of Notch signaling (*63, 65*).

### Micro-gavage assay

At 5 days post-fertilization (dpf), zebrafish larvae were manually dechorionated and immobilized in 1% low-melting-point agarose (ThermoFisher, MA, USA) containing neutralized tricaine (Sigma-Aldrich) for micro-gavage, as described previously (*64*). A gavage solution consisting of Rhodamine B isothiocyanate (R8881, Sigma-Aldrich) and 0.05% phenol red (Sigma-Aldrich) was delivered into the anterior intestinal bulbus using a microinjection system, taking care to avoid the esophagus, yolk sac, and swim bladder.

NPs or PA-MPs were suspended in the Rhodamine solution before administration. At 6 or 24 h post-gavage, larvae were imaged on a dual-channel confocal microscope (Leica SP8, Wetzlar, Germany) to assess Rhodamine disposition in the posterior cardinal vein (PCV) and caudal vein plexus (CVP).

### DNA constructs and Generation of transgenic line

A custom Tol2-based CRISPR/Cas9 construct (*pTol2-cmlc:EGFP-fli1ep:dsRed-Cas9*) targeting dre_piezo1 was generated by VectorBuilder (Chicago, USA) based on the method of Ablain *et al*. (*66*). This vector carries dual U6 promoters driving two *dre_piezo1* gRNAs (table S3), an endothelial-specific *fli1ep:Cas9-T2A-dsRed* cassette, and a *cmlc:EGFP* cardiac reporter for transgenic screening. Plasmid DNA (250 ng/ul) was co-injected with Tol2 transposase mRNA (250 ng/ul) into one-cell zebrafish embryos as previously described. Embryos expressing dsRed in endothelium were selected for further genotyping and phenotypic analyses. Scramble zebrafish gRNAs (Table 1) used as controls were retrieved from previous published sequences (*67, 68*).

### T7 Endonuclease I (T7E1) assay

Genomic DNA was isolated from individual 2 dpf zebrafish embryos using a simplified alkaline lysis method. Regions flanking the CRISPR/Cas9 target sites were amplified by PCR with primers (table S4) positioned approximately 150-200 bp on each side of the predicted cleavage site. Mutation detection was performed using the EnGen Mutation Detection Kit (NEB) following the manufacturer’s protocol. The digested fragments were separated on a 2% agarose gel, and genome editing events were confirmed by the appearance of cleavage bands.

### RNAscope analysis

Zebrafish larvae were fixed overnight in 10% neutral buffered formalin (NBF) at room temperature, rinsed in PBS containing 0.1% Tween-20 (PBST), and transferred to 100% methanol for storage at -20°C overnight. Following rehydration, larvae were washed 4 times in PBST (5 min each). Samples were treated with Protease Plus (ACDbio) for 30 min at 40°C, rinsed 3 times in PBST (10 min each), and hybridized with a *piezo1*-C1 probe (ACDbio) in a HybEZ Oven for 2 h at 40°C. Hybridization signals were amplified according to the manufacturer’s protocol (ACDbio). Larvae were stored in PBST at 4°C until confocal imaging (*69*).

#### *In vivo* Ca^2+^ test

To monitor the Ca^2+^ dynamics after PA-MPs exposure, we used *Tg(elavl3:GCaMP6s)* and *Tg(myl7:gCaMP4*.*1*^*LA2124*^) lines, which express the fluorescent Ca^2+^ sensor GCaMP in neuronal and cardiomyocytes, respectively. Larvae were anesthetized in 0.02% neutralized tricaine (Sigma-Aldrich) and mounted in 0.03% low-melting agarose (Sigma-Aldrich) on glass coverslips. Ca^2+^ signals were recorded using a fluorescence microscope (Olympus IX70). Representative images of *Tg(elavl3:GCaMP6s*) larvae were acquired using a spinning disk confocal microscope (Leica DMI6000) and analyzed with Napari. Brain Ca^2+^ imaging movies were exported from Fiji and encoded to MP4 format using ffmpeg. Cardiac Ca^2+^ imaging stacks (800 frames) were processed in the same manner to ensure consistent encoding parameters across tissues. For qualitative comparison, movies from each condition were arranged side-by-side with identical visualization settings (including scaling, labeling, and rendering parameters) across PBS, NPs, and PA-MPs groups.

### Locomotor activity measurement

Locomotor activity was assessed in zebrafish larvae at 6 dpf following exposure to PBS, NPs, or PA-MPs. Individual larvae were placed in separate wells of a 24-well plate (1 larva per well). Swimming behavior was recorded using a high-resolution digital camera (Amscope, #MU series 3.0MP). The total distance traveled and average swimming speed were quantified from video recording using the ImageJ water maze plugin (NIH) (*34*).

## Statistical analysis

All experiments were performed with three independent biological replicates unless otherwise stated. Statistical analyses were performed using GraphPad Prism (Version 10). Data are presented as mean ± SEM for normally distributed variables and as median with 95% confidence intervals for non-normally distributed data. Comparisons between two groups were made using an unpaired two-tailed Student’s t-test. One-way ANOVA was performed for comparisons among multiple groups, followed by Tukey’s post hoc test. A *P* value < 0.05 was considered statistically significant. Exact *P* values are reported in the figures.

## Acknowledgement

This project is supported by NIH 5T32HL144449 (J.M.C., E.Z., and T.H.) NIH R01 ES033660 (S.R. and T. H.), R01 ES032037 (E.F.C. and T.H.), R01 HL159970 (T.H.), and R01HL129727 (T.H.).

## Reference

1. W. Huang, X. Xia, Element cycling with micro(nano)plastics. Science 385, 933–935 (2024).

2. A. I. Osman et al., Microplastic sources, formation, toxicity and remediation: a review. Environ Chem Lett, 1–41 (2023).

3. A. Collins et al., Emerging investigator series: microplastic-based leachate formation under UV irradiation: the extent, characteristics, and mechanisms. Environ Sci (Camb) 9, 363–374 (2023).

4. R. C. Thompson et al., Twenty years of microplastic pollution research-what have we learned? Science 386, eadl2746 (2024).

5. F. Cheng, T. Zhang, Y. Liu, Y. Zhang, J. Qu, Non-Negligible Effects of UV Irradiation on Transformation and Environmental Risks of Microplastics in the Water Environment. J Xenobiot 12, 1–12 (2021).

6. T. Meng et al., Multiple coronary heart diseases are risk factors for mental health disorders: A mendelian randomization study. Heart Lung 66, 86–93 (2024).

7. P. Li et al., Direct entry of micro(nano)plastics into human blood circulatory system by intravenous infusion. iScience 26, 108454 (2023).

8. L. Geppner et al., A novel enzymatic method for isolation of plastic particles from human blood. Environ Toxicol Pharmacol 104, 104318 (2023).

9. L. C. Jenner et al., Detection of microplastics in human lung tissue using muFTIR spectroscopy. Sci Total Environ 831, 154907 (2022).

10. T. Horvatits et al., Microplastics detected in cirrhotic liver tissue. EBioMedicine 82, 104147 (2022).

11. S. Zhang, H. Chen, L. Li, Z. Li, D. Wang, Exposure to placental microplastic and placental and umbilical cord blood telomere length. Ecotoxicol Environ Saf 302, 118536 (2025).

12. R. Marfella et al., Microplastics and Nanoplastics in Atheromas and Cardiovascular Events. N Engl J Med 390, 900–910 (2024).

13. A. J. Nihart et al., Bioaccumulation of microplastics in decedent human brains. Nat Med 31, 1114–1119 (2025).

14. W. Wang et al., Effects of polyethylene microplastics on cell membranes: A combined study of experiments and molecular dynamics simulations. J Hazard Mater 429, 128323 (2022).

15. J. B. Fleury, V. A. Baulin, Microplastics destabilize lipid membranes by mechanical stretching. Proc Natl Acad Sci U S A 118, (2021).

16. P. J. Dalal, W. A. Muller, D. P. Sullivan, Endothelial Cell Calcium Signaling during Barrier Function and Inflammation. Am J Pathol 190, 535–542 (2020).

17. S. S. Ranade et al., Piezo1, a mechanically activated ion channel, is required for vascular development in mice. Proc Natl Acad Sci U S A 111, 10347–10352 (2014).

18. V. Caolo et al., Shear stress activates ADAM10 sheddase to regulate Notch1 via the Piezo1 force sensor in endothelial cells. Elife 9, (2020).

19. H. Zi et al., Piezo1-dependent regulation of pericyte proliferation by blood flow during brain vascular development. Cell Rep 43, 113652 (2024).

20. A. L. Duchemin, H. Vignes, J. Vermot, Mechanically activated piezo channels modulate outflow tract valve development through the Yap1 and Klf2-Notch signaling axis. Elife 8, (2019).

21. Y. Guo et al., Aggregation behavior of polystyrene nanoplastics: Role of surface functional groups and protein and electrolyte variation. Chemosphere 350, 140998 (2024).

22. Q. Li et al., Aged polystyrene microplastics exacerbate alopecia associated with tight junction injuries and apoptosis via oxidative stress pathway in skin. Environ Int 186, 108638 (2024).

23. H. Peace et al., Stable staining of microplastics using conjugated polymer nanoparticles. Environmental Science: Nano 12, 2229–2233 (2025).

24. R. Syeda et al., Chemical activation of the mechanotransduction channel Piezo1. Elife 4, (2015).

25. Q. Zhao et al., Structure and mechanogating mechanism of the Piezo1 channel. Nature 554, 487–492 (2018).

26. S. Yang et al., Membrane curvature governs the distribution of Piezo1 in live cells. Nat Commun 13, 7467 (2022).

27. W. J. Polacheck et al., A non-canonical Notch complex regulates adherens junctions and vascular barrier function. Nature 552, 258–262 (2017).

28. S. Gonzalez-Ramos et al., Integrating 4-D light-sheet fluorescence microscopy and genetic zebrafish system to investigate ambient pollutants-mediated toxicity. Sci Total Environ 902, 165947 (2023).

29. K. I. Baek et al., Vascular Injury in the Zebrafish Tail Modulates Blood Flow and Peak Wall Shear Stress to Restore Embryonic Circular Network. Front Cardiovasc Med 9, 841101 (2022).

30. H. Huang et al., Microplastics in the bloodstream can induce cerebral thrombosis by causing cell obstruction and lead to neurobehavioral abnormalities. Sci Adv 11, eadr8243 (2025).

31. Y. Zhou et al., Low-dose of polystyrene microplastics induce cardiotoxicity in mice and human-originated cardiac organoids. Environ Int 179, 108171 (2023).

32. N. Vladimirov et al., Light-sheet functional imaging in fictively behaving zebrafish. Nat Methods 11, 883–884 (2014).

33. H. Shimizu et al., Mitochondrial Ca(2+) uptake by the voltage-dependent anion channel 2 regulates cardiac rhythmicity. Elife 4, (2015).

34. Y. Liu, Y. Wang, N. Li, S. Jiang, Avobenzone and nanoplastics affect the development of zebrafish nervous system and retinal system and inhibit their locomotor behavior. Sci Total Environ 806, 150681 (2022).

35. J. Yuval, I. Langmore, D. Kochkov, S. Hoyer, Neural general circulation models for modeling precipitation. Sci Adv 12, eadv6891 (2026).

36. X. Li et al., UV/ozone induced physicochemical transformations of polystyrene nanoparticles and their aggregation tendency and kinetics with natural organic matter in aqueous systems. J Hazard Mater 433, 128790 (2022).

37. K. Zhu et al., Inorganic anions influenced the photoaging kinetics and mechanism of polystyrene microplastic under the simulated sunlight: Role of reactive radical species. Water Res 216, 118294 (2022).

38. L. Tong et al., Polystyrene microplastics sunlight-induce oxidative dissolution, chemical transformation and toxicity enhancement of silver nanoparticles. Sci Total Environ 827, 154180 (2022).

39. Y. Xu, Q. Ou, J. P. van der Hoek, G. Liu, K. M. Lompe, Photo-oxidation of Micro- and Nanoplastics: Physical, Chemical, and Biological Effects in Environments. Environ Sci Technol 58, 991–1009 (2024).

40. Y. Zhang et al., Enhanced toxic effects of photoaged microplastics on the trophoblast cells. Toxicol Lett 409, 32–41 (2025).

41. H. Cheng et al., Immunotoxicity responses to polystyrene nanoplastics and their related mechanisms in the liver of zebrafish (Danio rerio) larvae. Environ Int 161, 107128 (2022).

42. J. Zhao et al., Oral Polystyrene Consumption Potentiates Atherosclerotic Lesion Formation in ApoE(-/-) Mice. Circ Res 134, 1228–1230 (2024).

43. F. Wernig, Q. Xu, Mechanical stress-induced apoptosis in the cardiovascular system. Prog Biophys Mol Biol 78, 105–137 (2002).

44. S. Chien, Mechanotransduction and endothelial cell homeostasis: the wisdom of the cell. Am J Physiol Heart Circ Physiol 292, H1209–1224 (2007).

45. Z. Peng, B. Shu, Y. Zhang, M. Wang, Endothelial Response to Pathophysiological Stress. Arterioscler Thromb Vasc Biol 39, e233–e243 (2019).

46. H. Lee, S. J. Song, C. S. Kim, B. Park, Polystyrene nanoplastics-induced intestinal barrier disruption via inflammation and apoptosis in zebrafish larvae (Danio Rerio). Aquat Toxicol 274, 107027 (2024).

47. J. Yu, L. Chen, B. Wu, Size-specific effects of microplastics and lead on zebrafish. Chemosphere 337, 139383 (2023).

48. E. Chuntharpursat-Bon et al., PIEZO1 and PECAM1 interact at cell-cell junctions and partner in endothelial force sensing. Commun Biol 6, 358 (2023).

49. D. Douguet, A. Patel, A. Xu, P. M. Vanhoutte, E. Honore, Piezo Ion Channels in Cardiovascular Mechanobiology. Trends Pharmacol Sci 40, 956–970 (2019).

50. A. Lai et al., Analyzing the shear-induced sensitization of mechanosensitive ion channel Piezo-1 in human aortic endothelial cells. J Cell Physiol 236, 2976–2987 (2021).

51. E. E. Friedrich et al., Endothelial cell Piezo1 mediates pressure-induced lung vascular hyperpermeability via disruption of adherens junctions. Proc Natl Acad Sci U S A 116, 12980–12985 (2019).

52. S. Saleem, R. R. Kannan, Zebrafish: an emerging real-time model system to study Alzheimer’s disease and neurospecific drug discovery. Cell Death Discov 4, 45 (2018).

53. E. El Hayek et al., Photoaging of polystyrene microspheres causes oxidative alterations to surface physicochemistry and enhances airway epithelial toxicity. Toxicol Sci 193, 90–102 (2023).

54. E. Zhu et al., Bubble-Mediated Large-Scale Hierarchical Assembly of Ultrathin Pt Nanowire Network Monolayer at Gas/Liquid Interfaces. ACS Nano 17, 14152–14160 (2023).

55. E. Zhu et al., Biomimetic cell stimulation with a graphene oxide antigen-presenting platform for developing T cell-based therapies. Nat Nanotechnol 19, 1914–1922 (2024).

56. S. K. Park et al., Elevated arterial shear rate increases indexes of endothelial cell autophagy and nitric oxide synthase activation in humans. Am J Physiol Heart Circ Physiol 316, H106–H112 (2019).

57. J. M. Cho et al., Activating P2Y1 receptors improves function in arteries with repressed autophagy. Cardiovasc Res 119, 252–267 (2023).

58. F. Jiang et al., The mechanosensitive Piezo1 channel mediates heart mechano-chemo transduction. Nat Commun 12, 869 (2021).

59. A. Dobin et al., STAR: ultrafast universal RNA-seq aligner. Bioinformatics 29, 15–21 (2013).

60. F. A. Wolf, P. Angerer, F. J. Theis, SCANPY: large-scale single-cell gene expression data analysis. Genome Biol 19, 15 (2018).

61. B. Muzellec, M. Teleńczuk, V. Cabeli, M. Andreux, PyDESeq2: a python package for bulk RNA-seq differential expression analysis. Bioinformatics 39, (2023).

62. J. M. Cho et al., Habitual Exercise Modulates Neuroimmune Interaction to Mitigate Aortic Stiffness. Circ Res 136, 1579–1594 (2025).

63. J. Lee et al., 4-Dimensional light-sheet microscopy to elucidate shear stress modulation of cardiac trabeculation. J Clin Invest 126, 1679–1690 (2016).

64. K. I. Baek et al., An Embryonic Zebrafish Model to Screen Disruption of Gut-Vascular Barrier upon Exposure to Ambient Ultrafine Particles. Toxics 8, (2020).

65. J. J. Hsu et al., Contractile and hemodynamic forces coordinate Notch1b-mediated outflow tract valve formation. JCI Insight 5, (2019).

66. J. Ablain, E. M. Durand, S. Yang, Y. Zhou, L. I. Zon, A CRISPR/Cas9 vector system for tissue-specific gene disruption in zebrafish. Dev Cell 32, 756–764 (2015).

67. J. M. Uribe-Salazar et al., Evaluation of CRISPR gene-editing tools in zebrafish. BMC Genomics 23, 12 (2022).

68. F. Kroll et al., A simple and effective F0 knockout method for rapid screening of behaviour and other complex phenotypes. Elife 10, (2021).

69. J. Wang et al., Mechanically activated snai1b coordinates the initiation of myocardial delamination for trabeculation. Nat Commun 16, 8363 (2025).

