## Supplemental Figures for "Photoaged microplastics disrupt endothelial stretch-sensitive ion channels to impair calcium signaling and vascular integrity"

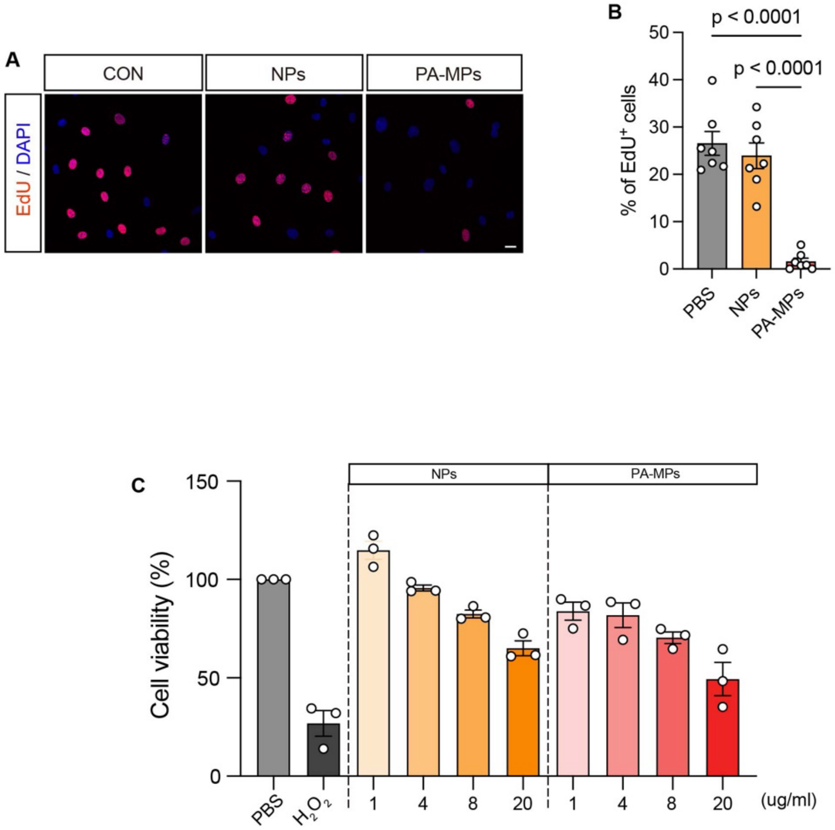


**Fig. S1. PA-MPs trigger attenuates endothelial growth.**

(A) Cell viability was assessed by the MTT assay after 24 h of exposure in a concentration-dependent manner, and cell proliferation was evaluated by EdU incorporation assay (B, C).

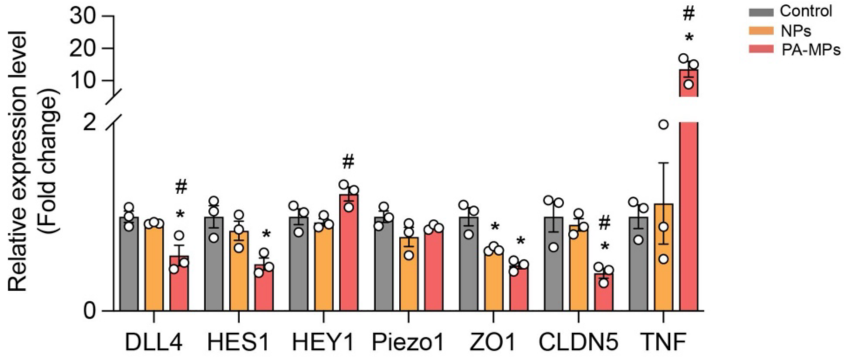


**Fig. S2. Independent qPCR validation corroborates RNA-seq-identified transcriptional changes.**

Quantitative PCR (qPCR) was performed to validate selected differentially expressed genes identified by bulk RNA-seq. Expression changes were consistent with sequencing results (n =3).

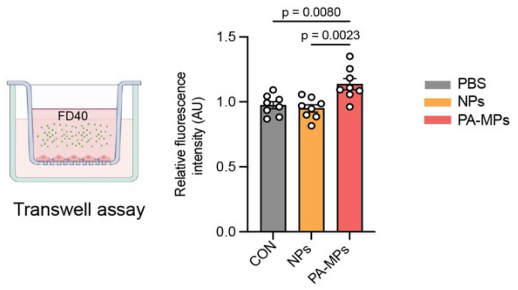


**Fig. S3. PA-MPs increase endothelial permeability.**
Transwell permeability assay revealed increased fluorescence intensity following PA-MPs exposure, indicating enhanced endothelial leakiness.

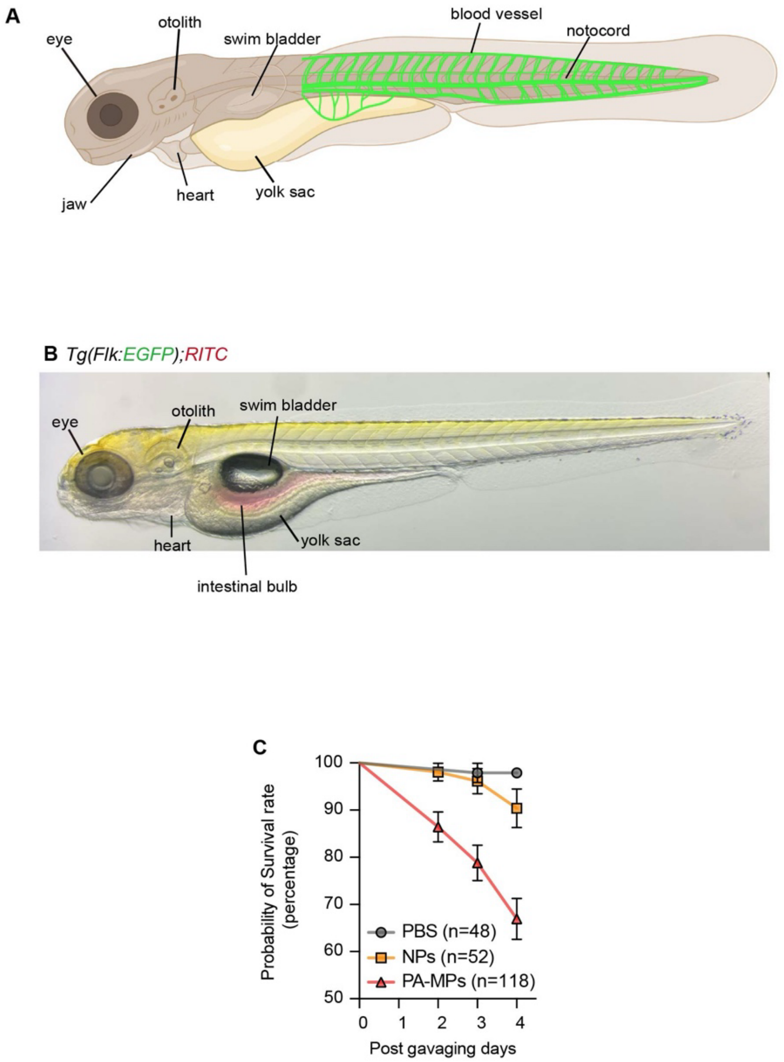


**Fig. S4. Verification of micro-gavage and survival analysis following PA-MPs exposure in zebrafish.**

(A, B) Representative image showing zebrafish larvae immediately after micro-gavage with microplastics mixed with Phenol Red, confirming accurate delivery into the intestinal lumen. (C) Survival curves of zebrafish from 0 to 4 days post-gavage (dpg) with PBS, NPs, or PA-MPs. The number of larvae assessed (n) is indicated next to each data point on the plot.

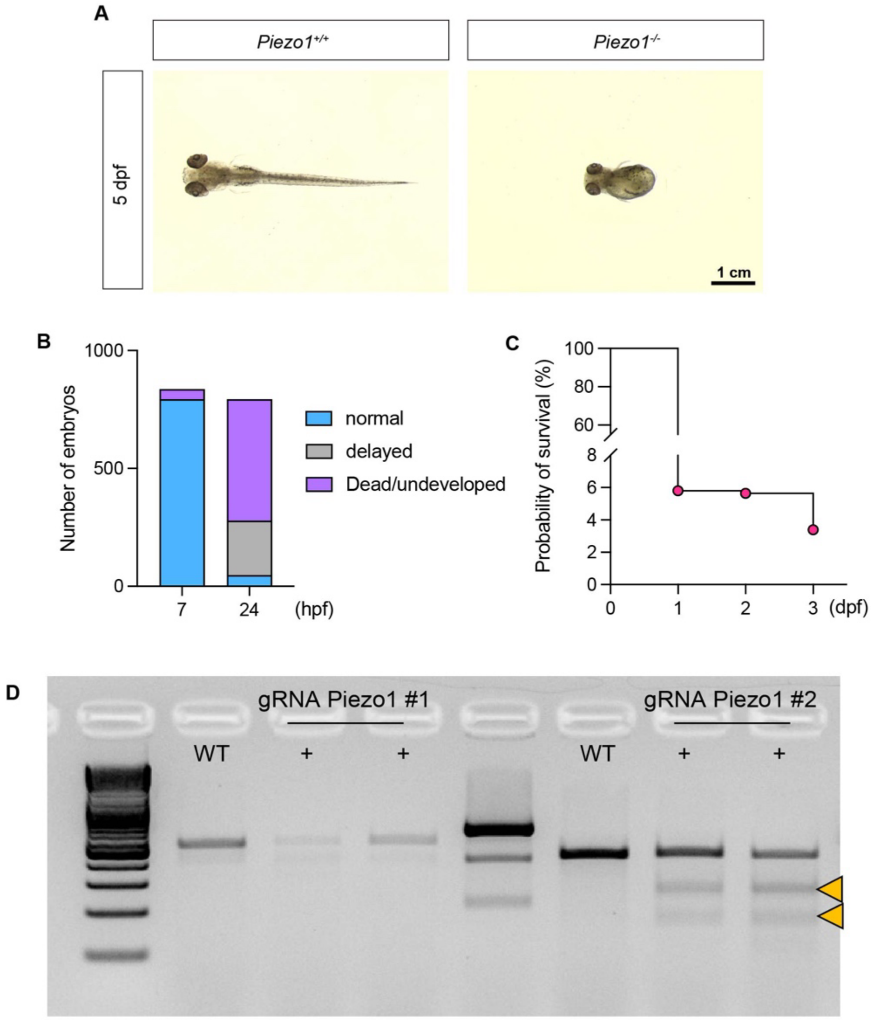


**Fig. S5. Validation of endothelial cell-specific *Piezo1* knockout zebrafish.**
(A) Representative images of *Piezo1* inbred zebrafish (wild type vs. endothelial-specific mutant). Loss of endothelial *Piezo1* expression impaired normal body development. (B) Classification of P1 embryos at 7 and 24 hours post-fertilization (hpf): blue, normal; gray, developmentally delayed; purple, dead or undeveloped. (C) Survival analysis showing a marked decline in viability beginning at 1 day post-fertilization (dpf) in the *Piezo1* mutant line. (D) T7E1 mutagenesis assay confirming CRISPR-Cas9-induced mutation at the piezo1 target site. Cleavage bands (arrowheads) indicate the presence of mutations at the target site. Genomic DNA collected from 8 embryos at 2 dpf showed distinct T7E1 cleavage products, indicated by arrows.

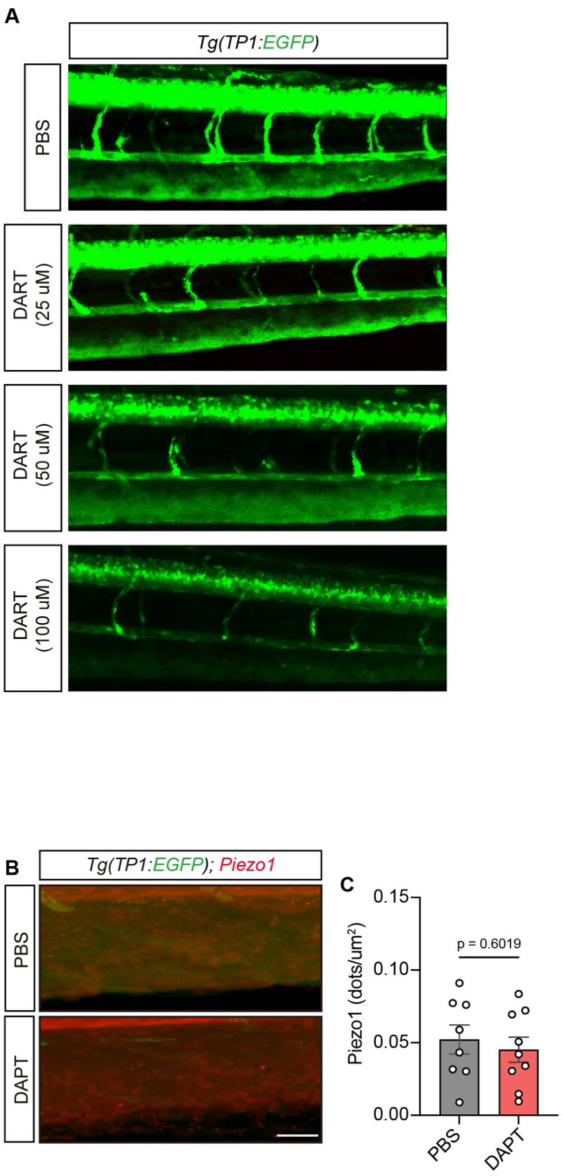


**Fig. S6. Verification of DAPT concentration for Notch inhibition in zebrafish.**
(A) To determine the optimal DAPT dose in *Tg(Tp1:EGFP)* zebrafish, we assessed Notch+ endothelial cells across a range of concentrations. Notch reporter activity decreased progressively with higher DAPT concentrations. From this assessment, 50 uM DAPT was identified as the optimal concentration for experimental use. (B, C) DAPT treatment, a γ-secretase inhibitor of Notch1 signaling, confirmed that Notch1 does not regulate *Piezo1* expression. n = 8-9 per group.

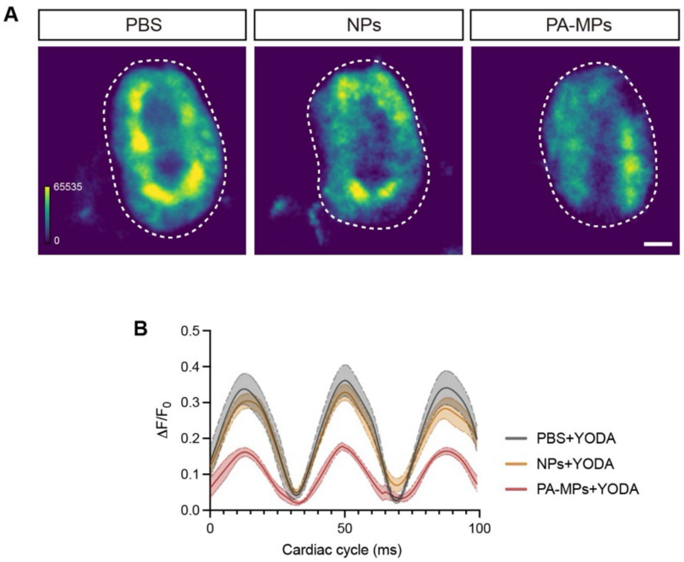


**Fig. S7. Yoda1 fails to restore calcium influx in PA-MPs-exposed zebrafish cardiomyocytes.** (A) Representative images of *Tg(myl7:gCaMP4.1$^{LA2124}$)* zebrafish larvae showing calcium transients in cardiomyocytes following 2 uM Yoda1 treatment for 5 minutes. (B) Quantification of calcium flux ($\Delta F/F_0$) demonstrates that Yoda1 markedly increased calcium signaling in PBS and NPs treated groups, whereas PA-MPs-exposed larvae showed no rescue of calcium influx. n = 5-10 per group.

**Table S1. List of antibodies for western blot analysis.**

| Protein | Supplier | Host | Dilution ratio | Clonality | Catalog No. |
|---|---|---|---|---|---|
| Piezo1 | Novus | Mouse | 1:500 | Monoclonal | NBP2-75617 |
| TRPV4 | Alomone Labs | Rabbit | 1:2000 | Polyclonal | ACC-124 |
| TRPC6 | Alomone Labs | Rabbit | 1:2000 | Polyclonal | ACC-120 |
| Notch1 | Cell Signaling | Rabbit | 1:1000 | Monoclonal | 4380 |
| Adam10 | Novus | Rabbit | 1:1000 | Polyclonal | NBP1-76973 |
| NICD | Cell Signaling | Rabbit | 1:1000 | Monoclonal | 4147 |
| PECAM | Abcam | Mouse | 1:800 | Monoclonal | ab24590 |
| VE-Cadherin | R&D systems | Goat | 1:800 | Polyclonal | AF938 |
| Occludin | Thermo Fisher | Rabbit | 1:250 | Polyclonal | 40-4700 |
| Claudin5 | Thermo Fisher | Mouse | 1:1000 | Monoclonal | 35-2500 |
| Cleaved Caspase-3 | Cell Signaling | Rabbit | 1:1000 | Polyclonal | 9661 |
| 4-HNE | Abcam | Rabbit | 1:1000 | Polyclonal | ab46545 |
| GAPDH | Proteintech | Mouse | 1:50000 | Monoclonal | 60004-1-lg |

**Table S2. List of primer sequences for qPCR.**

| Gene | Primer sequence (5’-3’) |
|---|---|
| DLL4 | Forward: ACAACTTGTCGGACTTCCAG |
| | Reverse: CAGCTCCTTCTTCTGGTTTG |
| ADAM10 | Forward: TTGCCTCCTCCTAAACCACTTCCA |
| | Reverse: AGGCAGTAGGAAGAACCAAGGCAA |
| HES1 | Forward: CCAAAGACAGCATCTGAGCA |
| | Reverse: GCCGCGAGCTATCTTTCTT |
| HEY1 | Forward: GCGTGGGAAAGGATGGTTGAG |
| | Reverse: TCCGCTCTCGGCTGCTTG |
| Piezo1 | Forward: CGTCTTCGTGGAGCAGATG |
| | Reverse: GCCCTTGACGGTGCATAC |
| ZO1 | Forward: CAACATACAGTGACGCTTCACA |
| | Reverse: CACTATTGACGTTTCCCCACTC |
| CLDN5 | Forward: CTCTGCTGGTTCGCCAACAT |
| | Reverse: CAGCTCGTACTTCTGCGACA |
| IFN | Forward: TCGGTAACTGACTTGAATGTCCA |
| | Reverse: TCGCTTCCCTGTTTTAGCTGC |
| TNF | Forward: CCTCTCTCTAATCAGCCCTCTG |
| | Reverse: GAGGACCTGGGAGTAGATGAG |
| GAPDH | Forward: GCCTCAAGATCATCAGCAAT |
| | Reverse: GGACTGTGGTCATGAGTCCT |

**Table S3. List of primer sequences for CRISRP-Cas9 system.**

| Gene | sgRNA sequence (5'-3') |
|---|---|
| Piezo1 sg #1 | TAAACCGTACAGCAGGATGA |
| Piezo1 sg #2 | ATGTTTGCGCTTGGGTGCCC |
| Scrambled sg #1 | CGTTAATCGCGTATAATACG |
| Scrambled sg #2 | CATATTGCGCGTATAGTCGC |

**Table S4. List of primer sequences for T7E1 assay.**

| Gene | sgRNA sequence (5'-3') |
|---|---|
| sgRNA1-Piezo1_Fw | TGATTCACGACCCTGAGAGC |
| sgRNA1-Piezo1_Rv | TGCATTTATTTTGAACCCAGAAAGT |
| sgRNA2-Piezo1_Fw | AGCTGGCTGACATTTGTGCT |
| sgRNA2-Piezo1_Rv | GCGATGTATTTTTAGAACAGCTTTG |
